# Approaches to measuring language lateralisation: an exploratory study comparing two fMRI methods and functional transcranial Doppler ultrasound

**DOI:** 10.1101/2023.09.01.555910

**Authors:** D. V.M. Bishop, Z. V. J. Woodhead, K. E. Watkins

## Abstract

In this exploratory study we compare and contrast two methods for deriving a laterality index (LI) from functional magnetic resonance imaging (fMRI) data: the weighted bootstrapped mean from the widely-used Laterality Toolbox (toolbox method), and a novel method that uses subtraction of activations from homologous regions in left and right hemisphere to give an array of difference scores (mirror method). Data came from 31 individuals who had been selected to include a high proportion of people with atypical laterality when tested with functional transcranial Doppler ultrasound (fTCD), and who had fMRI data for two tasks, word generation and semantic matching, which were selected as likely to activate different brain regions. In general, the mirror method gave better agreement with fTCD laterality than the toolbox method, both for individual regions of interest, and for a large region corresponding to the middle cerebral artery. LI estimates from this method had much smaller CIs than those from the toolbox method; with the mirror method, most participants were reliably lateralised to left or right, whereas with the toolbox method, a higher proportion were categorised as bilateral (i.e. the CI for the LI spanned zero). Reasons for discrepancies between fMRI methods are discussed: one issue is that the toolbox method averages the LI across a wide range of thresholds. Furthermore, examination of task-related t-statistic maps from the two hemispheres showed that language lateralisation is evident in regions characterised by deactivation, and so key information may be lost by ignoring voxel activations below zero, as is done with conventional estimates of the LI.

## Introduction

It has been established for many years that language processing depends on specialised areas in the left side of the brain in most people. Initial evidence came from observations of aphasia after brain lesions; as early as the mid 19th century it was noted that aphasia was strongly associated with left-sided damage (Berker et al., 1986). Neurosurgeons subsequently developed the Wada test (Wada & Rasmussen, 1960) in which each hemisphere was successively anaesthetised to determine which was dominant for language, again revealing the bias to left-sided language processing in most people. With the advent of modern brain imaging methods, it became possible to observe correlates of brain activation associated with different cognitive tasks, confirming the population bias to left hemisphere processing for generating language. There is, however, a significant minority of people who depart from the population trend, and have atypical lateralisation: this may take the form of reversal of the usual left-sided bias for language - right-hemisphere language - or an overall lack of bias - bilateral language.

Atypical lateralisation is of considerable theoretical interest to neuroscientists interested in the evolutionary origins and functional significance of laterality. In addition, it is relevant for understanding how individuals recover from unilateral brain injury and for planning epilepsy surgery.

The current study aims to advance progress in developing reliable, sensitive and valid methods for measuring functional brain lateralisation in individuals.

### Functional transcranial Doppler ultrasound (fTCD)

FTCD is an inexpensive and portable method for measuring lateralisation by comparing blood flow velocity in the left and right middle cerebral arteries (MCAs) during an activation task, relative to a resting baseline period. It is relatively insensitive to movement artefacts, and can be used with children aged 4 years and over (Bishop et al., 2014). In a recent study, Parker et al. (2022) compared lateralisation indices (LIs) for six different language tasks in left- and right-handers. Left-lateralisation characterised tasks involving language generation, but not tasks that predominantly involved receptive language, and lateralisation was generally weaker in left-handers. Split-half reliabilities were reasonable for all tasks (.7 or more).

With fTCD, signals from left and right sides are normalised by dividing by the mean blood flow velocity of that side, and each trial is baseline corrected by subtracting the mean value from a rest period prior to stimulus presentation. A sample plot from averaged data on word generation is shown in Figure 1, with the black line showing the (rescaled) difference between left and right channels. When fTCD was first developed as a tool for investigating laterality, the LI was obtained by identifying a temporal window, or Period of Interest (POI), and taking the mean value in an interval defined as +/- 2 s around the absolute peak difference between left and right signals in the POI (Deppe et al., 1997). This method had the disadvantage that it could give a biased LI estimate in weakly-lateralised individuals where the peak difference changed from left to right in the course of the POI. Selecting the larger peak could create a spurious bimodality in the distribution of LIs in a sample. As an alternative, Woodhead et al. (2020) proposed using a simple subtraction method whereby the mean signal in the right MCA was subtracted from that on the left. The two methods were highly correlated, but the subtraction method gave a normal distribution of LIs in a sample. More recently, Thompson et al. (2022) proposed a method more analogous to that used in fMRI, where Generalised Models were used to estimate the interaction of side (left vs right) with time interval (POI vs non-POI) on the signal over a series of trials. This gave LIs that agreed well with the simpler subtraction method, but with a smaller error of measurement. For the current paper, we use the complex Generalised Additive Model (GAM) method described by Thompson et al. (2022) to estimate LIs from fTCD data. This estimates the response function from the data, where predictors are side, POI (boxcar), plus terms for time relative to epoch start, plus a categorical index corresponding to individual epochs. LI is estimated as an interaction term: percentage difference in blood flow between left and right that is specific to the POI.

**Figure 1:**
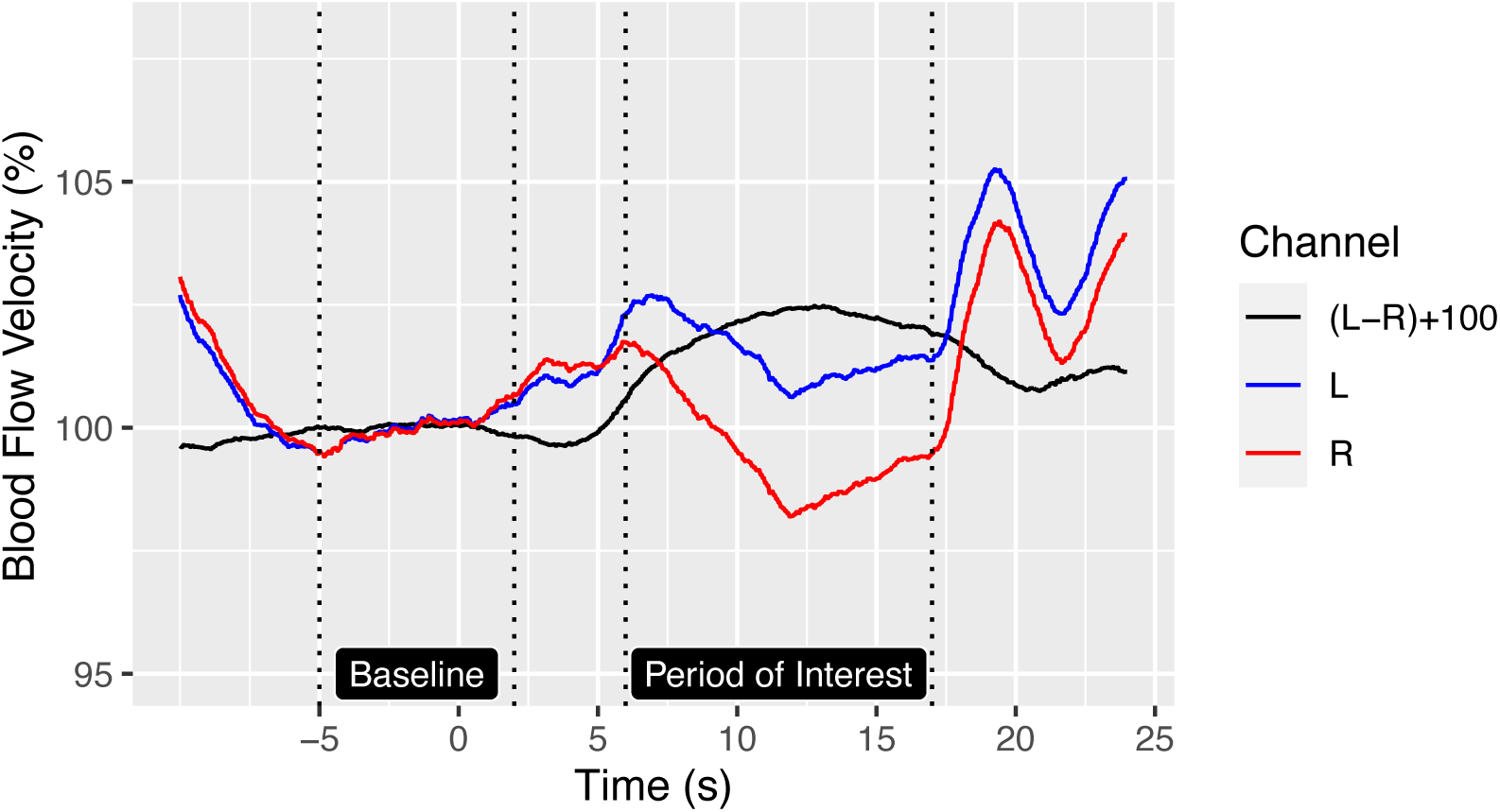
Averaged blood flow velocity over course of a word generation trial.

FTCD has excellent temporal resolution, but can only tell us about left-right differences, not where activations are occurring within a hemisphere. To answer questions about localisation, we need to turn to fMRI, which has complementary advantages and disadvantages to fTCD: it has poor temporal resolution but good spatial resolution, and can measure changes in blood oxygenation in voxels as small as 1 mm^3^. By comparing LIs obtained with fTCD (LI_fTCD_) and fMRI (LI_fMRI_) on comparable tasks, we can gain some insights into the sensitivity of fTCD to regional changes in blood flow. However, comparison is complicated by measurement issues.

### Functional Magnetic Resonance Imaging (fMRI)

A comprehensive review by Seghier (2008) described the range of methods that have been used to derive a LI from fMRI data. The optimal method for estimating laterality has been much debated (Bradshaw et al., 2017). Vingerhoets et al. (2023) used the Delphi approach (Hasson et al., 2000) to explore expert opinion on measurement of cerebral lateralisation. They concluded that there was still no agreement about the optimal way to measure lateralisation using fMRI, and the key concept of bilateral language was ill-defined. We need reliable and convenient methods for measuring functional lateralisation in individuals to advance research in this area. Here we compare a widely-used approach, the LI Toolbox, with an alternative approach, the mirror method.

### The LI-Toolbox

LI-toolbox is a freely available Matlab (Mathworks, Natick, MA, USA) toolbox that offers a flexible range of options for computing a laterality index (LI) (Wilke & Lidzba, 2007; Wilke & Schmithorst, 2006). The starting point is a t-statistic map derived in the conventional way from the General Linear Model, which is used to contrast the condition of interest with a comparison condition, which may be either rest or another condition that differs on key task demands. The next step is to identify voxels from the t-statistic map within the region-of-interest (ROI) that are above an activation threshold, and compute the LI by the standard formula (L-R)/(L+R) where L and R are either the number of voxels, or the summed activation of suprathreshold voxels on L and R sides. In practice, it makes little difference whether number of voxels or summed activation is used in the formula (Wilke & Lidzba, 2007). However, the choice of threshold can make a substantial difference to the LI. Figure 2 shows an illustrative example where the computed LI (based on summed t-values in activated voxels) rises from .33 to .75 between thresholds of zero to 6, and then declines, to become negative (i.e. right-biased) at thresholds of 12 or more.

**Figure 2:**
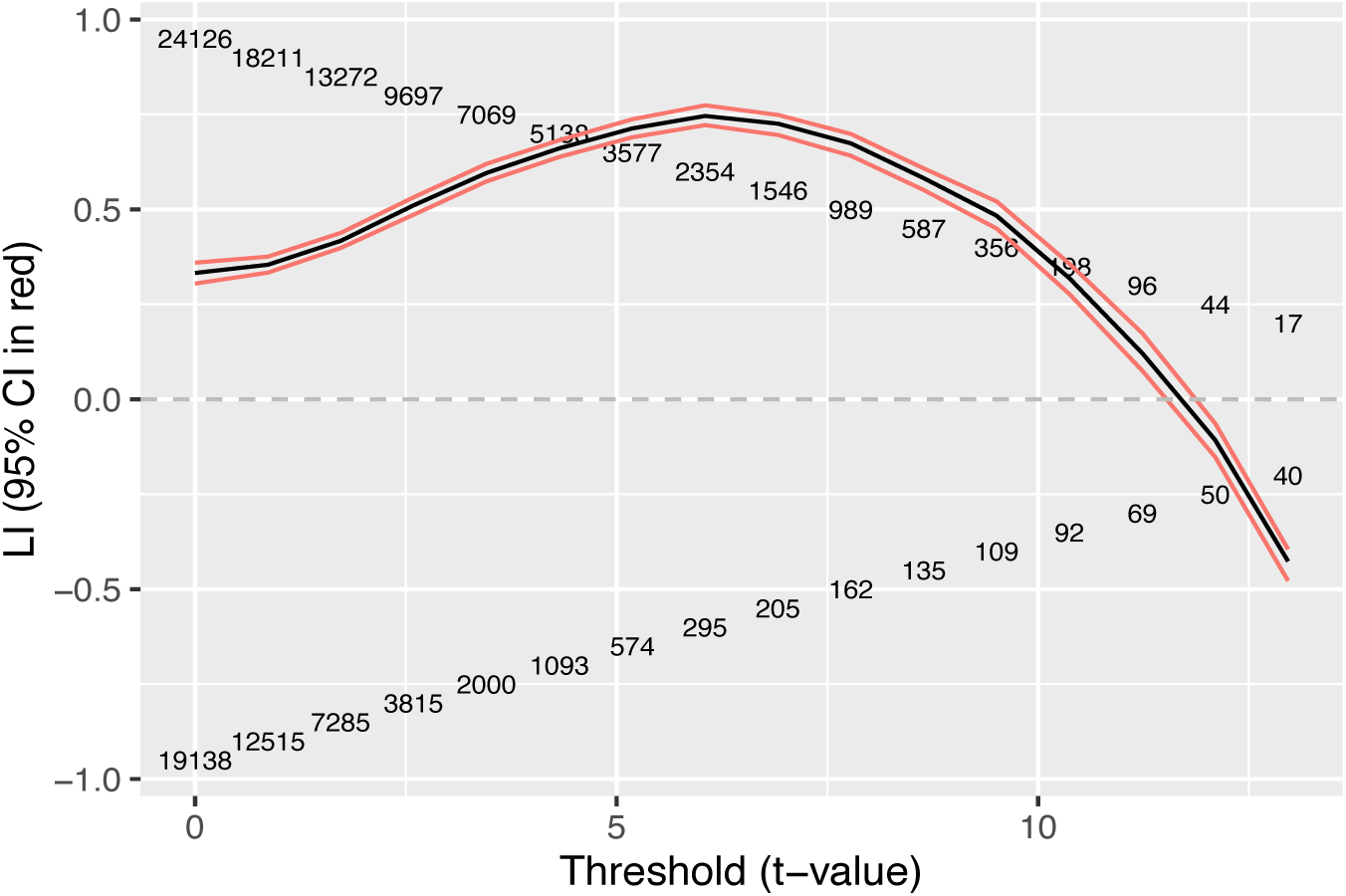
LI estimates (based on summed activation) at different thresholds in a sample individual fMRI dataset using a word generation task for activation. The numbers in the body of the graph show the numbers of suprathreshold voxels on the left (above zero) and right (below zero). There are 16 threshold t-values ranging from 0 to 12.96. Higher thresholds yielded too few voxels to meet criteria for inclusion.

One method from the LI toolbox that has become popular as a way of taking into account information from a wide range of thresholds involves bootstrapping: the paper by Wilke & Schmithorst (2006) describing this approach has received 185 citations in Web of Science at the time of writing. This method (described in Bradshaw et al., 2017; Wilke & Schmithorst, 2006) takes an iterative approach. First a set of 20 t-thresholds is created using equally-spaced steps from 0 to the maximum t-value in the image for the ROI. Threshold values resulting in fewer than 10 activated voxels on either side are discarded. At each remaining threshold, and for each hemisphere, the bootstrapping procedure starts by defining the set of suprathreshold voxels. From this set, 100 samples (with replacement) are taken of the t-statistics within the thresholded ROI; the default is to take 25% of values in each resample. So if in a given ROI there are 40 suprathreshold voxels in the left hemisphere and 24 in the right, 100 samples of 10 t-values are taken from the left, and 100 samples of 6 t-values from the right. An LI value is calculated based on the t-statistics for every left + right combination of these re-samples (100*100=10,000 LI values) at each threshold level. The bootstrapping approach has the advantage that a CI can be specified around the LI estimate for an individual. This makes it possible to have a statistical definition of bilateral language: rather than relying on an arbitrary cutoff, we can designate a person as having bilateral language if the 95% CI around the LI includes zero.

Sample data from the same participant as in Figure 2 are shown in Figure 3. Note that at any one threshold, the density plot of LIs is narrow, indicating little variation from iteration to iteration of the bootstrap, but the differences in LI between thresholds is substantial, and, as also shown in Figure 2, the LI for this participant becomes right-sided at very high thresholds.

**Figure 3:**
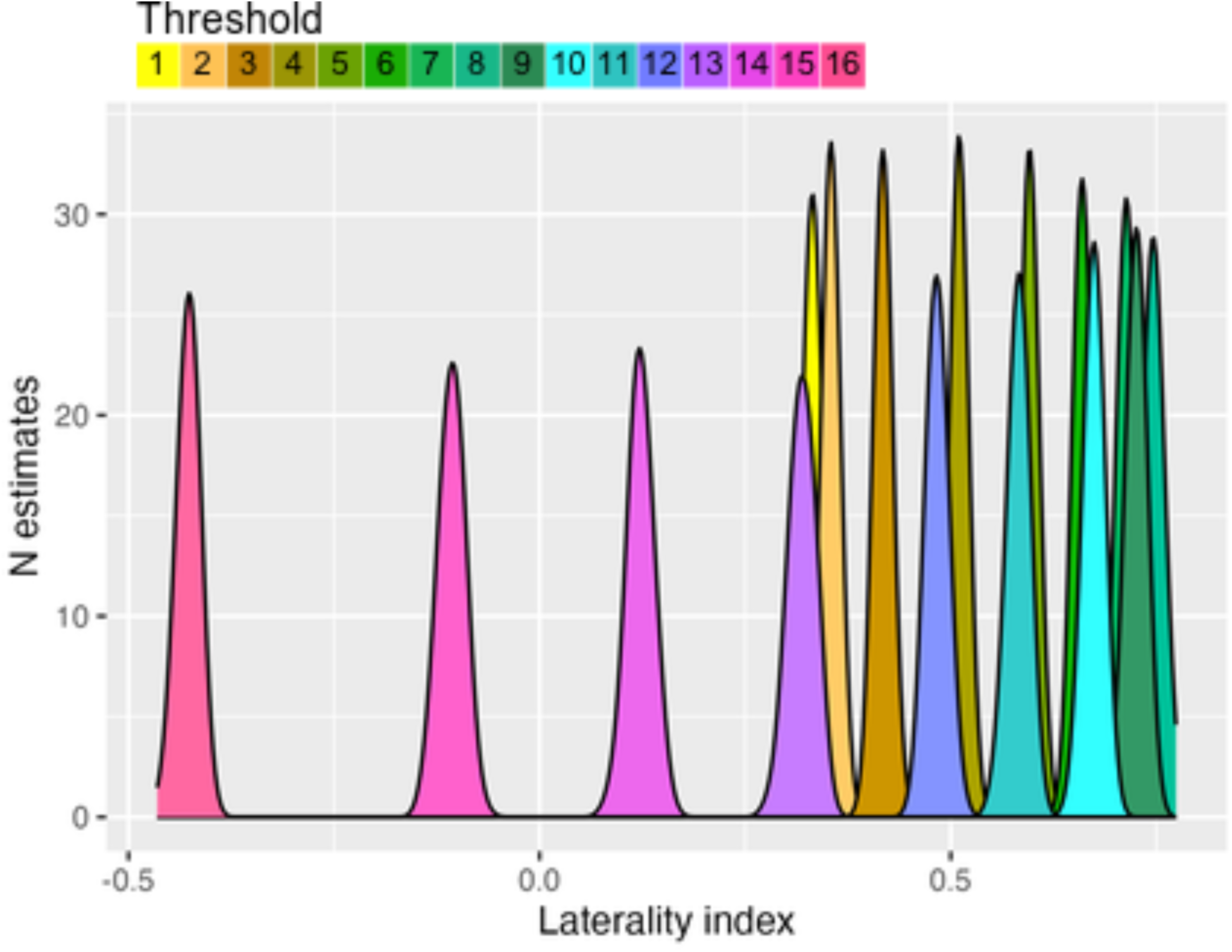
Distributions of bootstrapped LI estimates at 16 different thresholds in a sample individual fMRI dataset using a word generation task for activation. Threshold cutoffs are set in 20 steps from 0 to maximum activation: in this case the top four threshold cutoffs had insufficient voxels for inclusion

Alternative options are available for computing the LI from these estimates: The simplest is the mean of all the estimates - to reduce the impact of outliers, this is based on the middle 50% of values at each threshold, i.e., the 25% trimmed means. Further trimming can be conducted to give a trimmed mean based on the middle 50% of all of the trimmed mean LI values. Finally, one can compute a weighted mean that weights the trimmed mean LIs from each threshold by the threshold. Wilke & Schmithorst (2006) recommended this approach as it ensures that voxels with the highest correlation with the task will have most impact on the LI. All methods of mean estimation are intended to reduce the impact of outliers. Note that the 95% CI around these estimates is substantial; this does not indicate that the estimates are likely to be irreproducible. In fact, we know that the weighted mean LI from the LI toolbox has excellent test-retest reliability (Johnstone et al., 2020). The large CI reflects the fact that it is based on a wide distribution of thresholds; within any one threshold, the estimates of LI are very similar, but across the range of thresholds, the LI may vary substantially. Although the method is described as “threshold-free”, it is perhaps more accurate to say that it uses an algorithm for selecting a representative LI (either by mean, trimmed mean, or weighted mean), which will incorporate information from a wide range of thresholds.

Although the method is well-specified and justified, it still involves some arbitrary decisions. The bootstrap method is described as “threshold free”, but LI estimates still depend on the range of thresholds considered. This is evident from the wide CIs seen for LI estimates in Figure 4. These wide intervals do not mean that the computed LIs from the LI toolbox are not replicable, but rather that they are dependent on the specific range of thresholds that was selected. For instance, if we had taken a threshold range of t-values from 0 to 6, instead of 0 to 13 for the data in Figure 2, the LIs (mean, trimmed mean or weighted mean) would have been higher, the range of LI values lower, and the CIs substantially smaller. The LI-Toolbox uses a default threshold range extending to the maximum t-value observed in the image. It follows that the threshold range can vary from person to person, task to task, and ROI to ROI. Furthermore, fewer voxels are included at high thresholds, and so the least reliable LI estimates contribute most to the weighted mean. Another point is that the method excludes voxels with negative t-scores (i.e. below threshold of zero). It is generally assumed that only positive t-values indicating task-related activation are of interest; however, as will be discussed further below, for some tasks and brain regions, functional lateralisation is evident when there are negative t-values in both hemispheres, suggestive of task-related de-activation. These three considerations: range of thresholds, weighting by level of activation, and removal of voxels with negative t-values, are not necessarily problems for the method, but they do confirm that decisions about which settings to use require careful thought and will have consequences for LI estimates.

**Figure 4:**
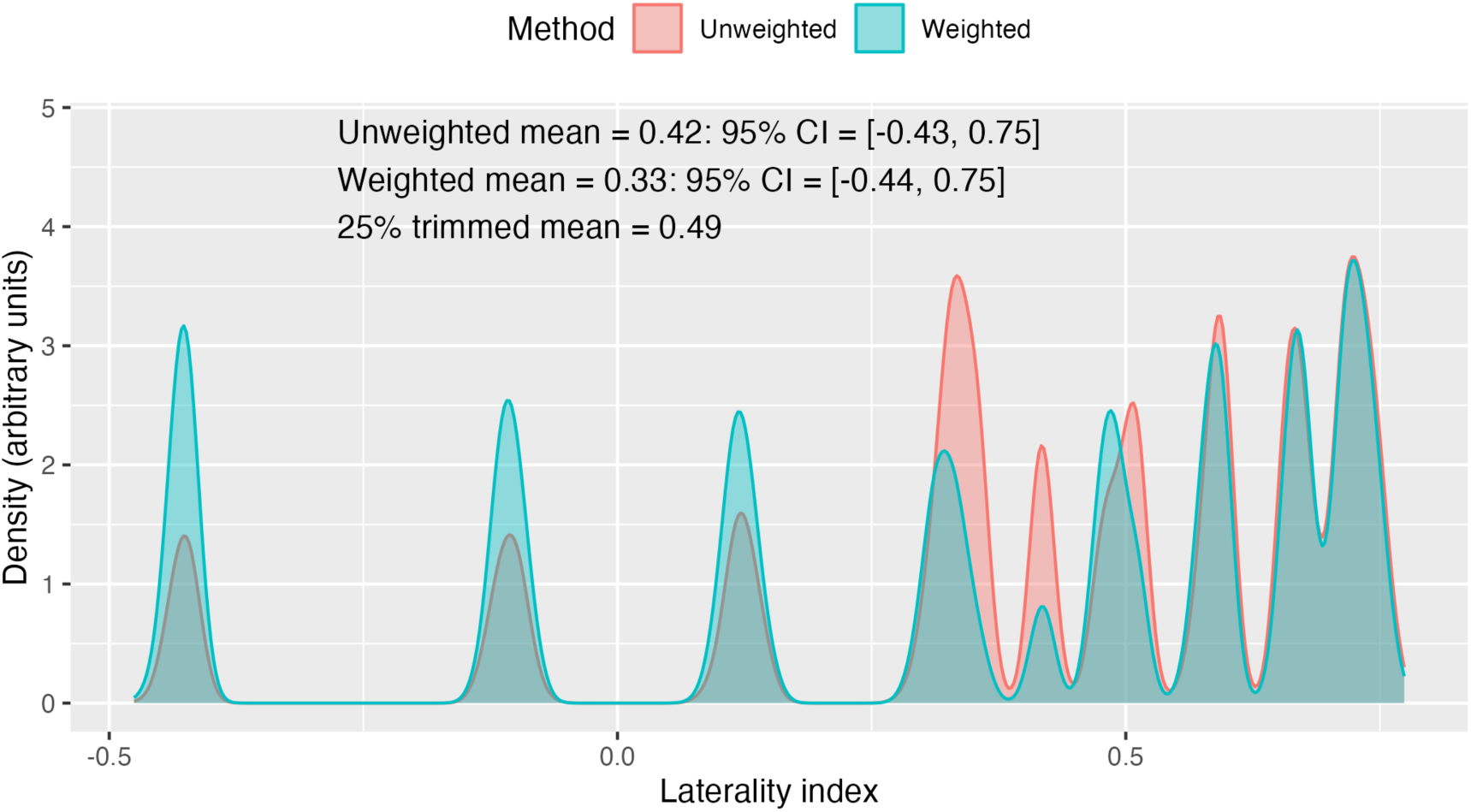
Weighted and unweighted distributions of bootstrapped LI estimates from the same dataset as Figure 2. The weighting makes the peaks at high thresholds higher, and those at low thresholds lower. The plot annotation shows LI estimates with different methods

### An alternative approach to fMRI laterality: The mirror method

We present here results from an alternative approach to measuring functional lateralisation, the mirror method, which was designed to be more comparable to the way in which laterality is assessed in fTCD. The logic is similar to the flip method, which was used by Watkins et al. (2001) to assess structural asymmetries in human MRI. In this method, rather than assessing how many voxels have suprathreshold t-statistics in left and right hemispheres, the right-brain t-statistic map is flipped on the left-right axis and then subtracted from the left-brain t-statistic map to give an map of a single hemisphere showing left-right differences in t-statistics for homologous regions.

This kind of method has occasionally been applied to functional MRI (Baciu et al.,2005; Cousin et al., 2007; Seghier et al., 2011), but there is no consensus on how to derive an individual LI using this approach. We term our method the mirror method to avoid confusion with other approaches that involve flipping one hemisphere before subtracting the activation maps.

Any method for determining laterality has to consider how to handle spatial dependencies between voxels. Smoothing and clustering are typically used (and options for clustering are provided in the LI Toolbox). Our approach with the mirror method was to minimise spatial dependencies between voxels within a region of interest by repeated sampling of a small set of values. Pilot testing showed that correlations between adjacent voxels in a sample were minimised when a sample of 5% voxels was used, and so 1000 samples (without replacement), each containing a random 5% of voxels in the region of interest, were taken, and the difference score computed at each voxel. The mean, 2.5th and 97.5th percentiles were taken from the 1000 samples to give the average difference score with a 95% CI. Note that this method is threshold-free, with all t-statistic map values being included, regardless of whether positive or negative.

### Prior comparisons of fMRI and fTCD

Thompson et al. (2022) compared estimates of LI from different analytic methods for a group of individuals who had completed similar language tasks with both fTCD and fMRI. A Generalised Additive Model (GAM) approach was developed to be more comparable to the Generalised Linear Model method used with fMRI. The individual LI values obtained with this method are closely comparable to those obtained by subtracting averaged left and right CBFV during the period of interest; however, the LI estimates have smaller standard errors than those from more traditional methods. The toolbox LI_fMRI_ was computed after applying a mask combining frontal, temporal and parietal lobes, as an approximation of the MCA territory. For the standard toolbox LI_fMRI_ (weighted mean from bootstrap), the correlation with LI_fTD_ was .66, indicating fair, but far from perfect agreement between methods. This raises the question of how far differences relate to method of computing the LI (difference score for fTCD and laterality index for fMRI), or better sensitivity to regional laterality effects in fMRI.

### Current aims

The main focus of Thompson et al. (2022) was on developing the GAM approach to analysing fTCD data. In contrast, our approach here is to consider in detail how LI_fTCD_ relates to LI_fMRI_, using the same dataset from Bruckert (2016). Following Seghier et al. (2011), we also aimed to see how laterality varied across different brain regions. We used data from two tasks, word generation and semantic matching, and four ROIs, frontal, temporal, parietal and cerebellar, and contrast the toolbox LI_fMRI_ and the mirror LI_fMRI_ with the LI_fTCD_.

As well as considering quantitative LIs, language laterality was categorised on each measure as left-lateralised, if the lower CI exceeded zero, right-lateralised if the upper CI was below zero, and bilateral if zero was included in the CI.

## Methods

### Design

A within-subjects design was used where participants were tested in two sessions: the first using fTCD, and the second using fMRI. On average, the interval between the two sessions was 11.8 days (range: 1 to 31 days).

In both sessions participants performed two tasks: a speech production task (word generation) and a semantic matching task based on the Pyramids and Palm Trees test (Howard & Patterson, 1992). An additional production task, auditory naming, was administered only in the fMRI session. The word generation and semantic matching tasks were designed to be as similar as possible in the two modalities (fTCD and fMRI), but some minor differences were required as noted below. Hence, the main within-subject independent variables were method (fTCD versus fMRI) and task (word generation versus semantic decision) and the dependent variable was laterality index (LI). For the semantic matching task, we administered a comparison (“baseline”) perceptual judgement task in the fMRI session, but here we present LIs from semantic matching vs rest, as this is comparable to the procedure used with fTCD. Further information about results with the perceptual judgement baseline task is provided in Supplementary Material 1. The auditory naming task was added to the analysis for comparisons of methods of laterality assessment in fMRI.

### Participants

Participants were recruited from a sample of 231 individuals who were screened with fTCD. These participants were recruited from the Oxford Psychology Research Participant Recruitment Scheme (https://opr.sona-systems.com). All participants gave written, informed consent, and all procedures were approved by the Central University Research Ethics Committee of the University of Oxford (MSD-IDREC-C1-20014-003). FTCD data from the full sample have been described by Bruckert et al. (2021).

In order to obtain a sample with cases of both typical and atypical lateralisation, all 21 participants who showed right lateralised or bilateral activation during word generation in fTCD were invited to return for the second session with fMRI. This classification was based on whether or not the 95% CI for the LI index on word generation crossed zero, where the CI was derived from the standard error of the LIs obtained on individual trials. Sixteen of these participants agreed to take part and fulfilled all MRI safety criteria. Sixteen participants with left lateralisation for word generation in fTCD, matched for age, gender and handedness, were also invited to return for the second session. The final sample that participated in both sessions comprised 32 participants (14 women, 18 men; 20 right handed, 12 left handed; mean age = 24.9 years, SD = 5.1 years). Due to time constraints, one participant missed the semantic matching task in the fMRI session, and another participant missed the auditory naming task, so the final sample size in the typical lateralisation group was 15 for those two tasks, and 16 for word generation.

### Functional Transcranial Doppler Ultrasound (fTCD)

#### Procedure

FTCD was acquired during the first session. Participants were trained to perform the two tasks and completed some practice trials; then the fTCD headset and probes were fitted and a stable signal was found. The fTCD signal was then recorded, first for the word generation task, then for the semantic matching task, with a break in between.

The timings for the two tasks are shown in Figure 5. Both tasks had a trial structure that started with a ‘Clear Mind’ instruction on screen (5 seconds), followed by the task for 20 seconds, and ending with 25 seconds of rest where the word ‘Relax’ was presented on screen. Participants were instructed to stay still and think of as little as possible during the ‘Relax’ period. The onset of the ‘Clear Mind’ cue was taken as the start of the trial (peri-stimulus time = 0s).

**Figure 5:**
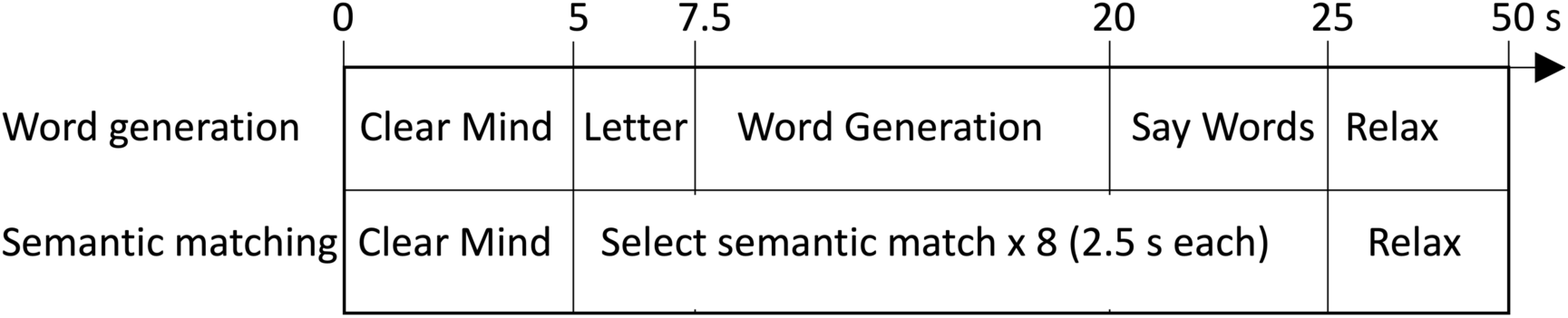
Time-course of trials in expressive (word generation) and receptive (semantic matching) tasks.

#### Word Generation

In this task (also known as verbal or phonemic fluency) a letter was presented in the centre of a screen and the participant was required to silently generate as many words as possible starting with that letter. There were 23 trials, each with a different letter (excluding Q, X and Z) presented in a randomised order. After the ‘Clear Mind’ cue, the letter was presented on screen for 2.5 seconds, followed by a blank screen for 12.5 seconds.

Participants were required to covertly generate words beginning with the letter during this period. A ‘Say Words’ cue was then presented for 5 seconds, during which participants were required to overtly report the words they had generated.

#### Semantic Matching

In the semantic matching task (based on the picture version of the Pyramids and Palm Trees test; Howard & Patterson, 1992) a triad of line drawings was presented, one at the top of the screen and two below. The participant was required to decide which of the two pictures below was the closest semantic match to the one at the top and respond by button press. There were 15 trials. For each trial, after the ‘Clear Mind’ cue, the participant was presented with eight consecutive picture triads, each lasting 2.5 seconds. Participants reported their decision by keyboard button press using their left or right index fingers. The location of the target picture was counterbalanced so that an equal number of left or right button presses was required. The time course of trials is shown in Figure 5.

#### Data Acquisition

The fTCD data were recorded from the left and right middle cerebral arteries (MCAs) simultaneously using two ultrasound monitoring probes held in place using an elastic headset. The signal was recorded using a Doppler-Box^TM^X receiver and processed using QL software (v3.2) on a laptop PC. All equipment (the probes, headset, receiver box and software) were from Compumedics DWL®. The experimental tasks were presented on a PC monitor using Presentation Software (Neurobehavioural Systems) which sent marker pulses to the Doppler-Box^TM^X system to denote the onset of each trial.

#### Data Analysis

The cerebral blood flow velocity (CBFV) data from left and right channels were analysed using custom scripts in R Studio (RStudio Team, 2015). The analysis followed the process described by Woodhead et al. (2019). In brief, this included downsampling from 100 Hz to 25 Hz; epoching from -12 s to 30 s peri-stimulus time; artefact rejection (both manual for gross artefacts and automatic for signal intensities outside of the 0.0001-0.9999 quantiles); signal normalisation; heart cycle integration; baseline correction using ten seconds of rest immediately preceding each trial as a baseline level; a final artefact detection stage where trials containing signal below 60% or above 100% of the mean normalised CBFV were rejected; and averaging of all trials (excluding rejected trials) for each task.

#### LI Calculation

For LI calculation we departed from the method used by Bruckert (2016) and adopted the complex Generalised Additive Model (GAM) approach described by Thompson et al. (2022), which is more comparable to the Generalised Linear Model method used with fMRI. The new LI_fTCD_ values from the GAM method were used to recategorise individuals as left-lateralised, right-lateralised, or bilateral on each of the two language tasks, depending on whether the 95% CI around the LI estimate was above zero (left), below zero (right), or spanned zero (bilateral).

The LI estimates obtained this way are not confined to the range -1 to 1. To make it possible to visualise LI_fTCD_ on the same scale as LI_fMRI_, values were scaled by dividing the LI_fTCD_ by 6; this meant that the the largest absolute scaled LI_fTCD_ value was close to 1.

#### Data Quality

Two prespecified criteria were used to check data quality. First, if any participant had more than 20% of trials rejected (i.e. more than five trials for word generation or more than three trials for semantic matching), they would be excluded from the analysis. Second, the trial-by-trial variability was assessed using the standard error of LI values for each trial. The fTCD data were previously analysed as part of a larger sample of 156 participants (Bruckert et al., 2021). Outlier standard error values were identified in that dataset, and excluded from the analysis. No participants were excluded from the current analysis on the basis of these two quality checks.

### Functional Magnetic Resonance Imaging (fMRI)

#### Procedure

FMRI was acquired in the second session. Participants were first briefed on the imaging protocol and practiced the tasks outside of the scanner. They were then positioned in the scanner and a structural brain image was acquired. A block design was used. Three tasks were performed in separate runs, each lasting six minutes. As well as word generation and semantic matching, an auditory naming task was used. This task had no fTCD counterpart, but is considered when looking at agreement between toolbox LI and mirror LI for different ROIs. The order of task presentation was counterbalanced between participants.

#### Word Generation

This task was similar to the fTCD version, but owing to fMRI’s greater susceptibility to motion artefacts there was no overt word reporting phase. A block design was used, where the task was performed for 15 seconds followed by 15 seconds of rest (with a fixation cross). There were 12 blocks, each with a different letter presented on the screen throughout the duration of the block. Participants were required to covertly think of as many words as they could starting with that letter. Twelve letters (A, B, E, G, J, K, M, N, O, S, U, V) were presented in a randomised order.

#### Semantic Matching

The presentation of picture triad stimuli was similar to the fTCD method. Each fMRI block comprised eight picture triads, each with a duration of 2.5 seconds (20 seconds in total). The participants were required to respond by button press using their left and right thumbs on an MRI-compatible button box.

Unlike the word generation task, a comparison task (active perceptual baseline) was also acquired during the semantic matching fMRI run. To maximise similarity with fTCD, this comparison task was not included in the current analysis, but further details can be found in Supplementary Material 1.

#### Auditory Naming

The auditory naming paradigm was based on the Auditory Responsive Naming task (Bookheimer et al., 1998) and identical to the one used by Badcock et al. (2012), who adapted this task for the use in fMRI. Participants heard short definitions of a high frequency nouns through MRI compatible in-ear headphones (model S14, Sensimetrics), and were required to silently generate the described word (e.g. the participant heard ‘shines in the sky’ and thinks of ‘sun’). Because this task used an auditory presentation, which creates substantial activation in auditory cortex, a reversed speech condition (same recordings played backwards) was included, so the effect of auditory stimulation could be controlled for.

#### Data Acquisition

Scanning was performed in a Siemens 3T Trio scanner with a 32-channel head coil. The task stimuli were presented using Presentation Software (Neurobehavioural Systems) with stimulus onset synchronised with the scanner. The stimuli were projected via a mirror mounted on the head coil.

A high resolution T1-weighted MPRAGE was acquired for image registration (TR = 2040 ms, TE = 4.7 ms, flip angle = 8°, 192 transverse slices, 1 mm isotropic voxels). Echo-planar images were acquired to measure change in blood oxygen levels during the behavioural tasks (TR = 3s, TE = 30 ms, flip angle = 90°, 48 axial slices, slice thickness = 3 mm, in-plane resolution = 3 x 3 mm).

#### Data Analysis

Data analysis was conducted using FEAT (the fMRI Expert Analysis Tool) in FSL (FMRIB Software Library, http://www.fmrib.ox.ac.uk/fsl). The preprocessing stages included head motion correction through realignment to the middle volume of the EPI dataset; skull stripping using FSL’s Brain Extraction Tool (BET; Smith, 2002); spatial smoothing using a 6 mm full-width-half-maximum Gaussian kernel; high-pass temporal filtering at 90 seconds; and unwarping using fieldmaps in FSL’s Phase Region Expanding Labeller for Unwrapping Discrete Estimates tool (PRELUDE) and FMRIB’s Utility for Geometrically Unwarping EPI (FUGUE; Jenkinson, 2003).

The preprocessed data were entered into first-level (subject-specific) general linear models (GLMs). The different task runs were analysed separately. The explanatory variables (EVs) in the GLM were: the timecourse of the active tasks convolved with a double-gamma function; the temporal derivatives of the timecourse EV; and six motion correction parameters as covariates of no interest. For word generation and semantic matching tasks, the contrast of interest was the implicit (resting) baseline. For auditory naming, the contrast of interest was the backward speech condition. FSL FLIRT (Jenkinson et al., 2002) was used to linearly transform t-statistic maps into standard space using the MNI152_T1_2mm_brain template.

#### LI Calculation

The following methods for calculating LI_fMRI_ were compared:

1. Conventional weighted mean LI from LI Toolbox

The t-statistic maps were used to calculate LI values using the bootstrapping method from the LI Toolbox (Wilke & Lidzba, 2007; Wilke & Schmithorst, 2006) for each task and ROI. Here we used the weighted mean, as described above, which we refer to as the toolbox LI.

The LI Toolbox software does not provide a CI for the weighted mean. We computed one by creating a new histogram of LIs, with the proportion of LIs weighted by threshold (see Figure 4 for an example). The mean of these values agrees with the weighted mean computed by the LI toolbox. The 2.5th and 97.5th centiles were determined for each weighted histogram to give a 95% CI.

2. Mirror method

As described above, the right-brain t-statistic map is flipped and then subtracted from the left-brain t-statistic map, and then 1000 samples, each with 5% of voxels, are sampled without replacement from the difference map. The mean difference score is the LI, and the 2.5th and 97.5th percentiles from the 1000 estimates give a 95% CI around the LI. The MNI152 T1 2mm brain template is not perfectly symmetric: 0.53% of voxels on the right have no homolog on the left, and 0.93% of voxels on the left have no homolog on the right. These voxels are ignored when the difference score is computed.

To make it easier to visualise the mirror method data on the same scale as the LI toolbox data and the LI_fTCD_, the differences were divided by the absolute maximum difference in the sample, 2.7, so that the scale maximum was 1.0. This is a linear transform so does not affect correlations. Note, however, that these values are not conventional laterality indices, as they are simple differences, rather than proportions. We refer to these as scaled mirror LI.

#### Regions of Interest

Initial comparisons of methods were done using a combined mask of the frontal, temporal and parietal lobes as an approximation of the MCA territory. This mask was selected for comparability to the fTCD method, where the LI is based on comparisons of blood flow in left and right MCAs.

The left and right mean activations and LI values were then calculated separately for frontal, temporal, parietal and cerebellar ROIs, as defined by masks in the LI Toolbox, which are based on a population-based atlas (Hammers et al., 2003; Wilke & Lidzba, 2007). Note that we anticipate that cerebellar lateralisation will be opposite to that seen for other ROIs (Jansen et al., 2006; Seghier et al., 2011).

In all analyses a region 5 mm either side of the midline was excluded from consideration, again using a template provided in the LI Toolbox.

### Data Analysis

We first present averaged t-statistic maps showing fMRI patterns of activation for the three tasks, averaged for those who were selected as left-, bilateral or right-lateralised (based on fTCD word generation). We also present t-statistic maps used in the mirror method, i.e. maps of the L-R t-values obtained by subtraction of the flipped right hemisphere values from the homologous regions of the left hemisphere.

The next set of analyses was conducted to look at the agreement between LI estimates from fMRI and those from fTCD for word generation and semantic matching. The similarity between LI_fMRI_ and LI_fTCD_ was assessed using scatterplots and bivariate correlations. As we anticipated that the LI values may not fit a normal distribution, we used Spearman’s rank correlations instead of Pearson’s correlations. Bootstrapping was used to calculate the 95% CIs around the Spearman correlations (Hervé, 2022).

Finally, we consider agreement between the three tasks and four ROIs for the two methods of computing LI_fMRI_.

## Results

Patterns of activation from t-statistic maps are shown in Figure 6 for averaged data from laterality subgroups (left, bilateral and right) for the three tasks, imaged with MRIcroGL (https://www.nitrc.org). Note that transparent coloured areas correspond to subcortical activations. These maps clearly show left-sided lateralisation in frontal regions for the left-lateralised group, which is most pronounced for word generation and auditory naming. A more mixed picture is seen for the bilateral and right-lateralised groups. It is clear from these maps that positive activation in language regions with production tasks is accompanied by deactivation (negative t-values) in the parietal region.

**Figure 6:**
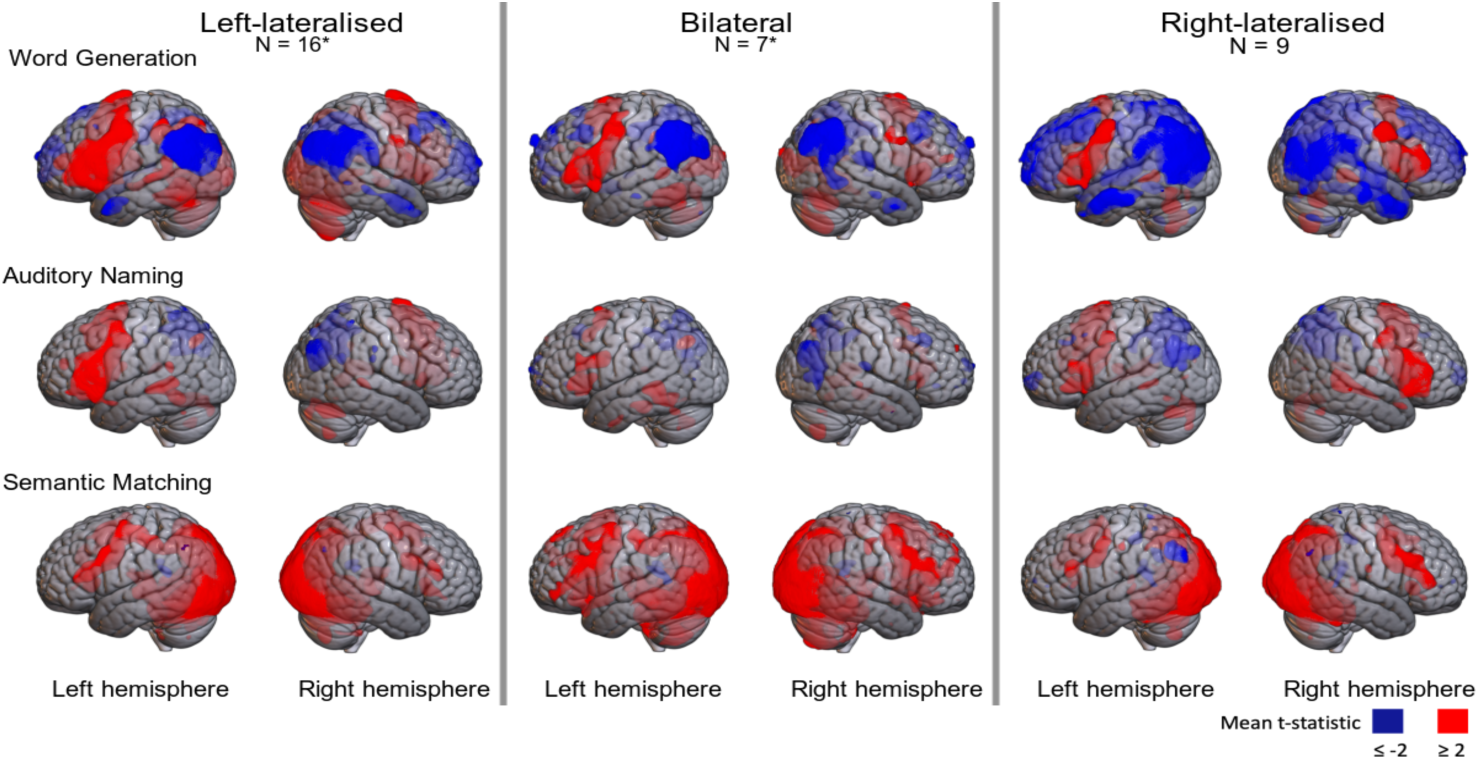
Heatmaps of t-statistic maps for the three tasks superimposed on SPM152 brain; averages for group defined by fTCD laterality on Word Generation. *Missing data reduces N by one for Auditory Naming bilateral group, and for Semantic Matching, left-lateralised group.

Figure 7 shows the subtracted t-statistic maps used in the mirror method, with lateralisation group based on fTCD word generation. Here, green indicates left-lateralisation, and yellow right-lateralisation. The left-lateralised group shows both frontal and posterior left-lateralisation (green) on word generation and auditory naming; note that the posterior left-lateralisation reflects the fact that negative t-values are seen in both left and right parietal regions in these tasks, but the negativity is more extreme in the right hemisphere, so the difference score favours the left side. The bilateral group also appears left-lateralised on these tasks, but with lateralisation confined to smaller regions. The right-lateralised group shows right-lateralisation (yellow) in frontal and posterior regions, but again lateralisation is less extensive than for the typically lateralised group. Note, however, that the averages for the bilateral and right-lateralised groups are based on smaller samples. These maps also emphasise the lack of lateralisation on semantic matching in the left-lateralised and bilateral groups.

**Figure 7:**
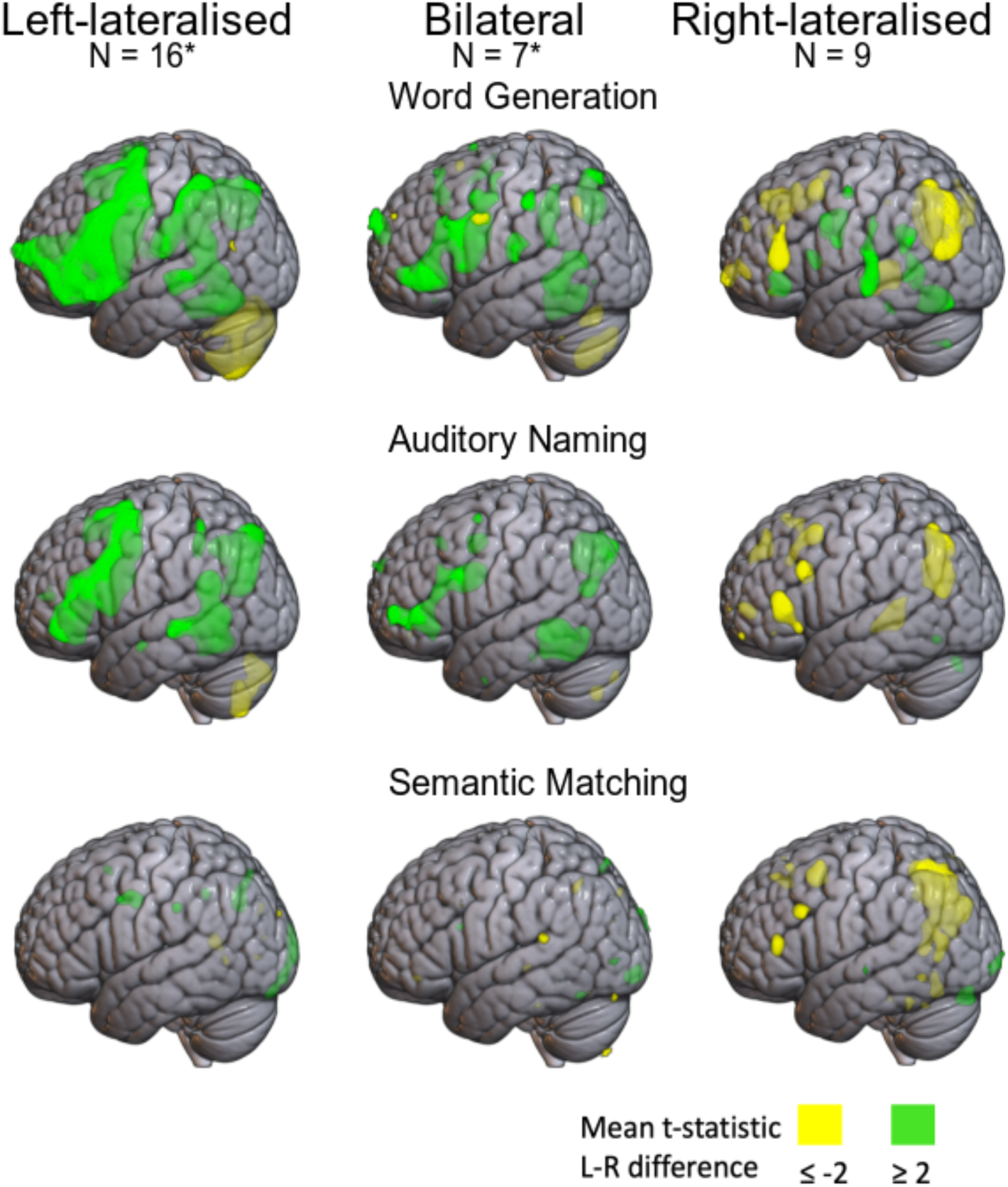
Heatmaps of difference t-statistic maps used in the mirror method for the three tasks superimposed on SPM152 brain; averages for left-, bilateral, and right-lateralised groups (defined by fTCD Word Generation). Green denotes left-lateralised voxels, and yellow denotes right-lateralised voxels. *Missing data reduces N by one for Auditory Naming bilateral group, and for Semantic Matching, left-lateralised group.

### 2. Correspondence of LI in the MCA from fTCD and fMRI, according to task and fMRI-LI method

Figure 8 shows the relationship between fTCD laterality (x-axis) and fMRI laterality (y-axis) for the original LI toolbox bootstrap method in the left panels and the new mirror method in the right panels. The upper panels show results for word generation, and the lower panels for semantic matching. Points are colour-coded according to categorical laterality classification on LI_fTCD_ for that task: grey points are those where the 95% CI crosses zero, and are hence coded bilateral. Points in red are right-lateralised, and those in blue are left-lateralised. Note that the high proportion of bilateral/right-lateralised cases reflects the fact that the sample was selected to include 50% atypically lateralised individuals. The plot also shows the Spearman correlation between fTCD and fMRI laterality, with the 95% CI. The percentage of individuals identified as having bilateral language on fMRI is also shown, again using the criterion of whether the 95% CI of the estimate crosses zero. Not shown in the plots is the correlation between laterality estimates from the two fMRI methods: the Spearman correlation [95% CI] between mirror and toolbox methods was 0.759 [0.526, 0.871] for word generation and 0.858 [0.685, 0.939] for semantic matching.

**Figure 8:**
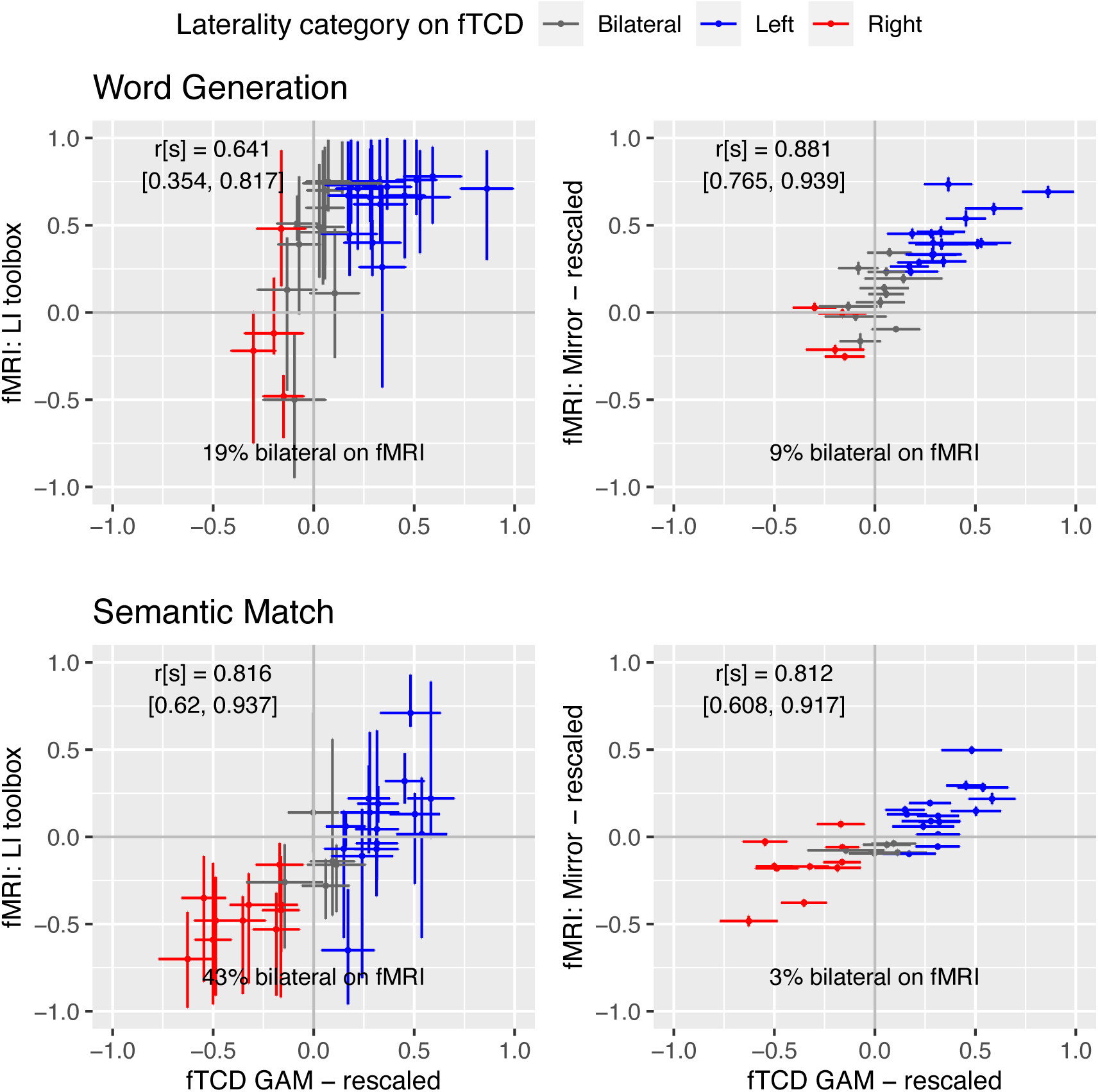
Scatterplots of relationship between fTCD and two methods of fMRI laterality for MCA region. Spearman correlations with 95% CIs are shown as annotations.

Two points are immediately evident from the plots. First, with few exceptions, fMRI and fTCD categorisations agree fairly well, i.e., those categorised as right-lateralised on fTCD tend to have LI values below zero on fMRI, and those categorised as left-lateralised on fTCD tend to have LI values above zero on fMRI. Second, the 95% CIs for individual LI estimates are considerably larger for the bootstrapped LI toolbox estimates than for the mirror method: this has the consequence that a far higher proportion of toolbox estimates categorise the individual as having bilateral language.

We will discuss possible explanations of these results below, but first we turn to look at agreement between LIs for the four ROIs outlined above. (See Supplementary Material 2 for details of the mean voxel activations on left and right for each region that are the basis for the LI calculations.)

Figure 9 shows correspondence of each fMRI laterality method with fTCD LI for each ROI for Word Generation. In general, the correlation is stronger for the mirror LI_fMRI_ than for toolbox LI_fMRI_, with the absolute value of the mirror LI_fMRI_ correlation being outside the 95% CI of that for the toolbox LI_fMRI_ correlation for all but the temporal ROI. The difference is most striking for the parietal region. The correlations with fTCD laterality are positive, except for the cerebellar region, where a negative association is seen.

**Figure 9:**
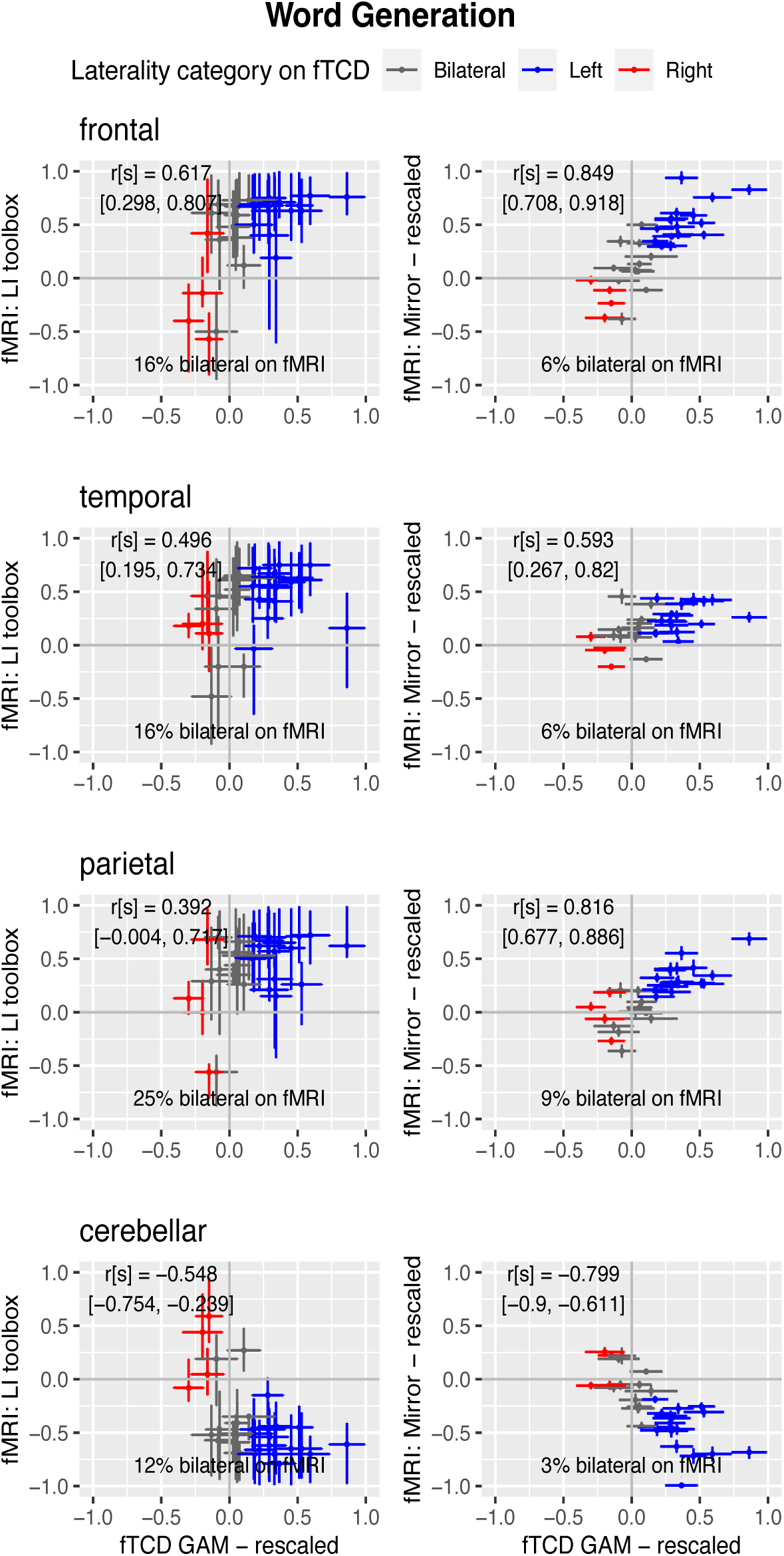
Scatterplots of relationship for Word Generation between fTCD and two methods of fMRI laterality for 4 ROIs

Figure 10 shows correspondence of each fMRI laterality method with fTCD LI for each ROI for Semantic Matching. The pattern of correlations is more similar for the two methods for this task, and it is noteworthy that for the cerebellum, the CIs for the correlations with LI_fTCD_ include zero for both measures.

**Figure 10:**
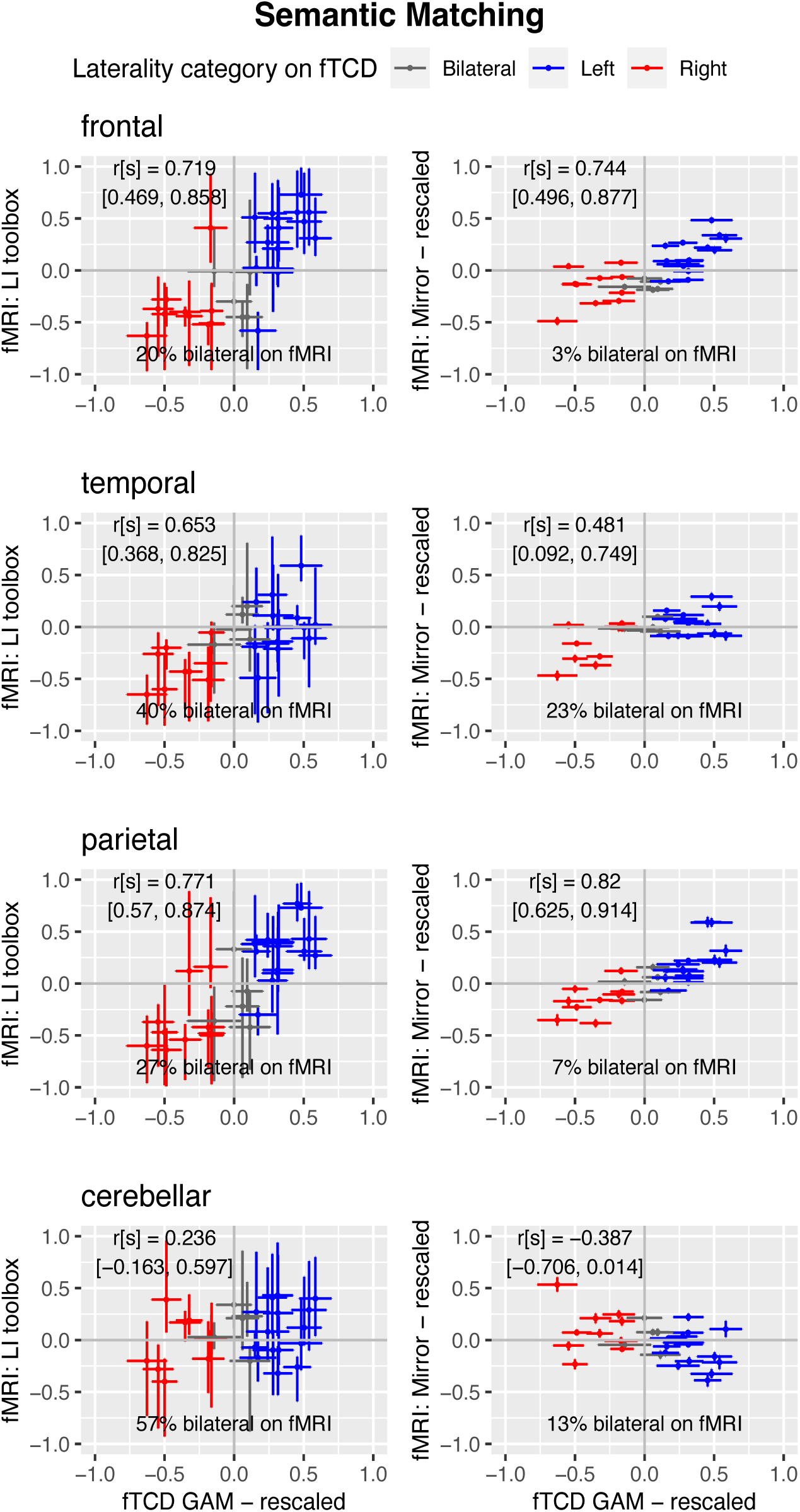
Scatterplots of relationship for Semantic Matching between fTCD and two methods of fMRI laterality for 4 ROIs

For both tasks, as was seen with the MCA ROI, the proportion of individuals categorised with bilateral language is much smaller with the mirror method than with the toolbox method, reflecting the large CIs associated with the toolbox estimates. In other words, people who appear to have bilateral language with the toolbox method appear reliably lateralised to left or right with the mirror method.

### 3. Consistency of laterality across tasks and ROIs

Spearman correlations between LIs from the three tasks and four ROIs are shown in Figure 11. Considering first the within-task correlations for the mirror method, one can see that for all three tasks, there is a positive correlation between frontal and parietal LIs, and a negative correlation between frontal/parietal and cerebellar LIs. Correlations with temporal regions are generally similar in direction to frontal correlations, but weaker. In addition, LIs for a given ROI are generally positive across the three tasks. Similar trends are seen for the toolbox LIs, but correlations tend to be a bit smaller in magnitude.

**Figure 11:**
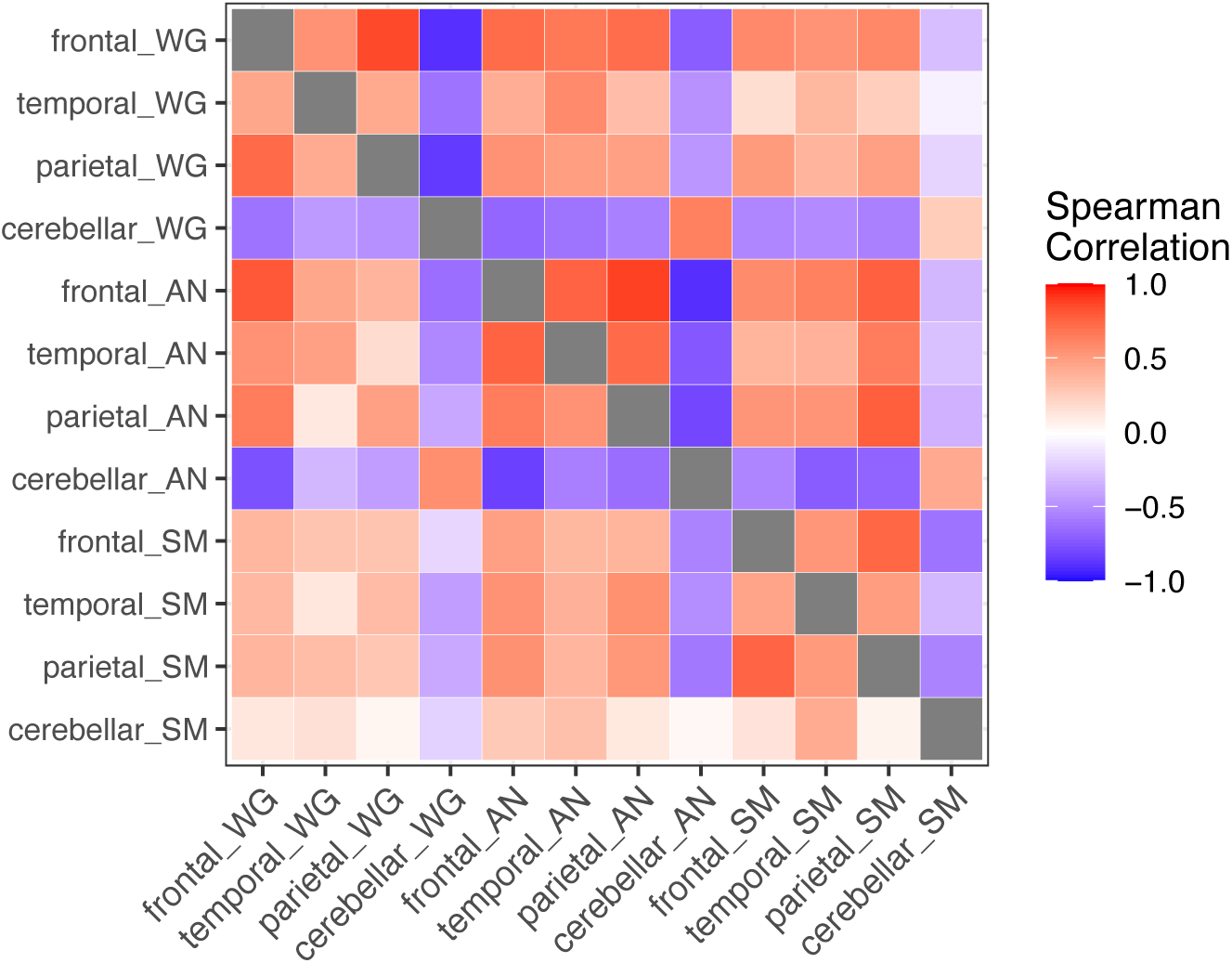
Heatmap of Spearman correlations between LIs for three tasks and four ROIs. Mirror method is above the diagonal, and toolbox method below the diagonal. Tasks denoted by WG (word generation), AN (auditory naming) and SM (semantic matching)

## Discussion

Estimates of LI_fMRI_ using the mirror method generally agree well with LI_fTCD_, whereas the toolbox LI has somewhat weaker correlations. It is important to note that the LI_fTCD_ cannot be regarded as a gold standard, as it does not have any sensitivity to localisation of activation. One possible explanation for the better agreement with mirror LI_fMRI_ is that the data processing is more similar to that in fTCD, involving simple subtraction of measures of activation on the left and right.

There is, however, another reason why the toolbox LI may give weaker agreement with LI_fTCD_: like virtually all methods of computing LI from fMRI, the toolbox LI_fMRI_ ignores task-related decreases in activation. This is justified by a simple assumption, stated by Wilke & Lidzba (2007) as: “‘interesting’ data will be of above-average intensity”.

### Importance of task-related deactivation

Figure 6 suggests that language laterality may depend on deactivation as well as activation. Indeed, this is already suggested by scrutiny of the mean task-related blood flow in the fTCD data for the word generation task (Figure 1). We first see an increase in blood flow to both hemispheres as the participant starts to perform the task, but then blood flow declines in both hemispheres, with the right hemisphere values dropping below the baseline level.

Our fMRI results have some similarities with those of Seghier et al. (2011), who considered how a global LI related to activation of individual voxels in left and right hemispheres.

Using a written semantic matching task, they found negative correlations between global LI and activations in some right hemisphere regions - i.e. the stronger the overall left-lateralisation, the lower the right-sided parameter values. This was not discussed in terms of deactivation, but could reflect a similar phenomenon as we report here. Task-related deactivation has been described for language tasks by Jansen et al. (2006), who noted that it poses problems for calculation of a conventional LI, because it can lead to values greater than 1. For instance, if mean activation is 2 for a left-ROI and is -1 for a right-ROI, then the computed LI is 3/1 = 3. If left- and right-ROIs have similar absolute activations opposite in polarity, the LI approaches infinity. Quite simply, the conventional LI is a proportional measure that is not appropriate for negative values. This means that we either have to ignore deactivation, or abandon the use of the conventional LI.

We suggest that deactivation appears to be an important feature in the brain’s lateralised response. The brain maps in Figure 6 show that deactivation is particularly striking in the parietal lobe. The quantitative analyses of average left and right activation in different ROIs in Supplementary material 2 confirm that, even for those with typical language lateralisation, we see average decrease of activation below zero in both left and right hemispheres, with the deactivation being more marked on the right. Furthermore, as can be seen in Figure 11, there are positive correlations between LI in the frontal and parietal ROIs, suggesting a common mechanism determines the extent of lateralisation. There is also a negative correlation with the frontal/parietal LIs and that obtained from the cerebellar ROI; this is evident in all three tasks when the mirror method is used.

The mechanism underlying such phenomena is not understood, though we know that the circle of Willis allows for compensatory changes in blood flow between posterior and anterior cerebral blood vessels, and between the two sides of the brain (Payne, 2017). It is possible that task-evoked neural activity that results in increased flow of oxygenated blood to the left frontal lobe will lead to a corresponding decrease in blood flow in posterior regions, and in the right hemisphere relative to the left.

If, as seems to be the case, functional lateralisation for language is manifest in deactivation of some regions, then we will fail to detect this phenomenon by relying on a laterality index that focuses only on voxels with suprathreshold activation.

### Categorisation of bilateral language

Another difference between the two methods for computing LI_fMRI_ is in the size of CIs around LI estimates, which are larger for the toolbox LI. It follows that if bilateral language is categorised using the CI around the mean estimate, then a high proportion of cases will be classed as bilateral using the toolbox method. The CIs around the mirror LI estimates are surprisingly small, given that they were based on relatively small samples of voxels in the difference map within a region. If we look only at averaged maps such as in Figure 6, we may get the impression that those with atypical lateralisation have similar activation on left and right sides of the brain during language tasks. The mirror LI estimates suggest that averaged maps may obscure the fact that most individuals have a significant bias to left or right-sided language, even for a weakly lateralised task such as semantic matching.

Development of the LI Toolbox was a major breakthrough in laterality measurement from fMRI, providing a serious attempt to tackle the inconsistent and arbitrary methods that were being used to quantify individual differences. The method allows one to compute a fast and reliable measure of LI_fMRI_ that has been shown to be robust to outliers, and is supported by open and thoroughly-documented code. This makes it straightforward to explore how different settings of parameters can influence the LI estimates. The wide CIs obtained with toolbox estimates do not reflect instability of the measures: rather, they reflect the fact that when a large set of possible thresholds is considered, the range of LI_fMRI_ estimates compatible with the observed data is large. This forces us back to the question of what is the best criterion for selecting a threshold, and we run into a difficult choice: a weighted mean is useful for giving most weight to the strongest evidence, but at the same time it means we rely most heavily on the least reliable estimates, based on a small number of voxels. In addition, if we take either the trimmed or weighted mean value from the distribution of LIs, this will be influenced by the range of t-values used to compute thresholds. Ultimately, while it is good to have an agreed convention of how to compute the LI_fMRI_, there remains some arbitrariness in how this is done.

Other studies have compared different approaches to estimation of laterality from fMRI data (see, e.g., Fesl et al., 2010; Abbott et al., 2010; Brumer et al., 2020), but, as far as we are aware, they all use the classic LI formula, (L-R)/(L+R), varying in how ROI or thresholds are determined, and whether L and R are number of activated voxels, or summed activation on left and right; in general, therefore, they do not take deactivation into account.

### Future Directions

In the course of comparing left- and right-sided activation associated with language generation, we noted that differences between hemispheres are not always driven by positive activation; in parietal lobes in particular, left-sided language processing tended to reflect reduced activity in both hemispheres, which is more pronounced on the right side. An unexpected benefit of our novel method, the mirror method, is that it allows us to capture this source of asymmetry without needing to specify any thresholds. This could be useful in future when comparing extent of lateralisation across different tasks or different regions of interest, and might cast light on the underlying mechanisms of functional lateralisation.

We have focused largely on estimating parameters and have presented few statistical tests here because we lacked *a priori* hypotheses and with 32 participants had limited statistical power to detect differences between methods, ROIs and tasks. This study may be regarded as hypothesis-generating rather than hypothesis-testing, and may be used as the basis for making predictions in future analyses of new datasets. In particular, we propose that:

*Prediction 1*: The pattern of activation and deactivation of left and right hemispheres in different regions seen in Figure 6 will replicate, with tasks involving language generation showing strong left hemisphere bias in all regions, but with a gradient from overall activation in frontal regions to overall deactivation in parietal regions.

*Prediction 2*: Cerebellar activation will show a complementary pattern of lateralisation to frontal activation for tasks involving language generation, but not for receptive language tasks.

*Prediction 3*: Most individuals who are categorised as having bilateral language on conventional fMRI laterality indices will show significant bias to left or right when assessed using the mirror method. In other words, most cases of bilateral language are probably cases of weak lateralisation which is hard to detect because of measurement error, rather than equal involvement of both hemispheres.

In addition, the results obtained here raise questions about the anatomical basis of individual differences in language lateralisation, suggesting that in future there might be value in relating functional differences to anatomical differences in the circle of Willis (Payne, 2017; Kızılgöz et al., 2022).

## Data Availability

Locations of data and analysis scripts are specified on Open Science Framework: https://osf.io/f6uyt/?view_only=23e92add2cbf4b1b9bf8a4816014e9b0 (Anonymous link for peer review).

## Acknowledgements

We are most grateful to Lisa Bruckert for making data from her doctoral thesis available to us for this study, and to Marko Wilke for helpful discussions about the LI Toolbox. This study was supported by H2020 European Research Council Advanced Grant 694189 (CANDICE).

