## Supplementary material 1 and 2 for "Approaches to measuring language lateralisation: an exploratory study comparing two fMRI methods and functional transcranial Doppler ultrasound"

### Semantic matching task with baseline task subtraction

The visual presentation of pictures in the semantic matching task leads to substantial activation of visual cortex. In the fMRI sessions (but not fTCD) a comparison condition was included as a control for the visual stimulation. In this perceptual condition, triads of abstract line drawings were presented in the same format as for the semantic matching task. Participants were required to detect which of the two drawings at the bottom was a perceptual match to the target drawing at the top and respond by button press.

The run comprised six blocks of semantic matching, six blocks of line decision and six rest blocks (where participants saw a fixation cross for 20 s), presented in a pseudo-randomised order where no condition was shown twice in a row.

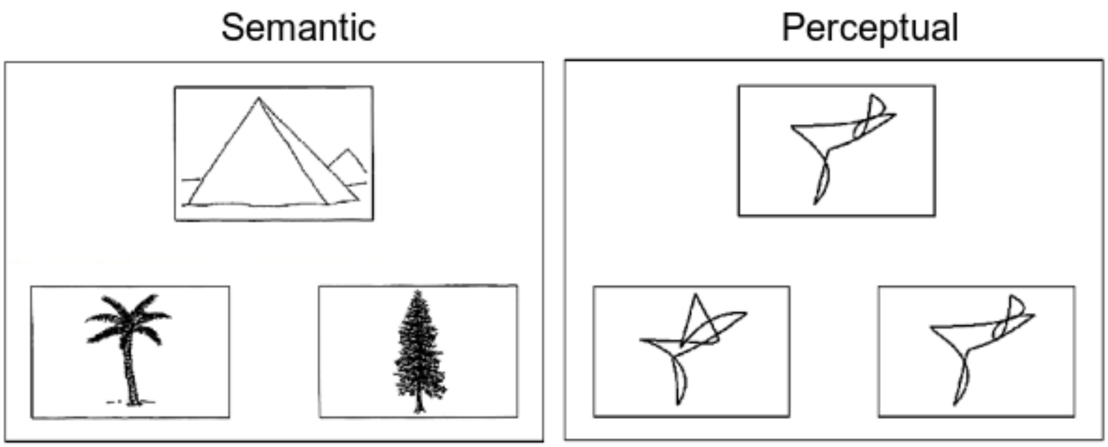


*Figure S1: Stimuli for semantic (left) and perceptual (right) matching conditions in the 'pyramids and palm trees' task*

For comparability to the fTCD task, our main analysis of fMRI data focused on the semantic matching task vs. rest, without any subtraction of the perceptual task activation. For completeness, we show here the patterns of activation for the individual semantic matching, perceptual matching, and contrast between those conditions (semantic minus perceptual).

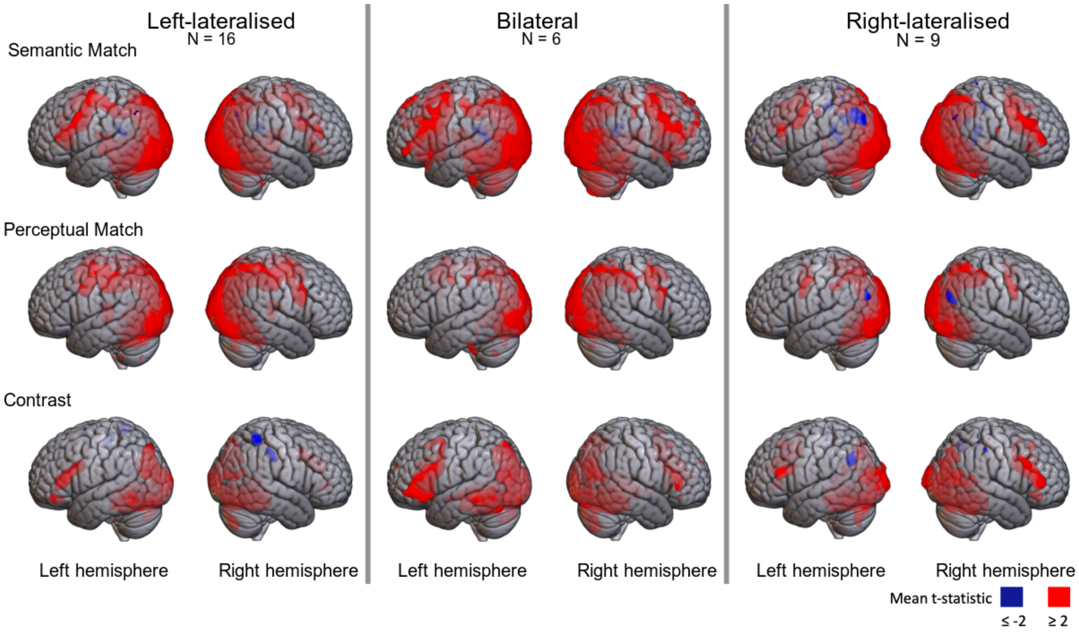


*Figure S2: Patterns of brain activation on the semantic and perceptual 'pyramids and palm trees' conditions, and the contrast between them for groups according to lateralisation category on fTCD Word Generation*

Focusing first on the typical group, one can see that there is frontal activation in both Semantic and Perceptual conditions, but it is more extensive on the left side in the Semantic task, as clearly seen in the Contrast condition. For the atypical group, frontal activation is similar on both left and right sides in the semantic task (see Contrast condition). These impressions are confirmed by the maps of L-R differences in t-values, shown in Figure S4.

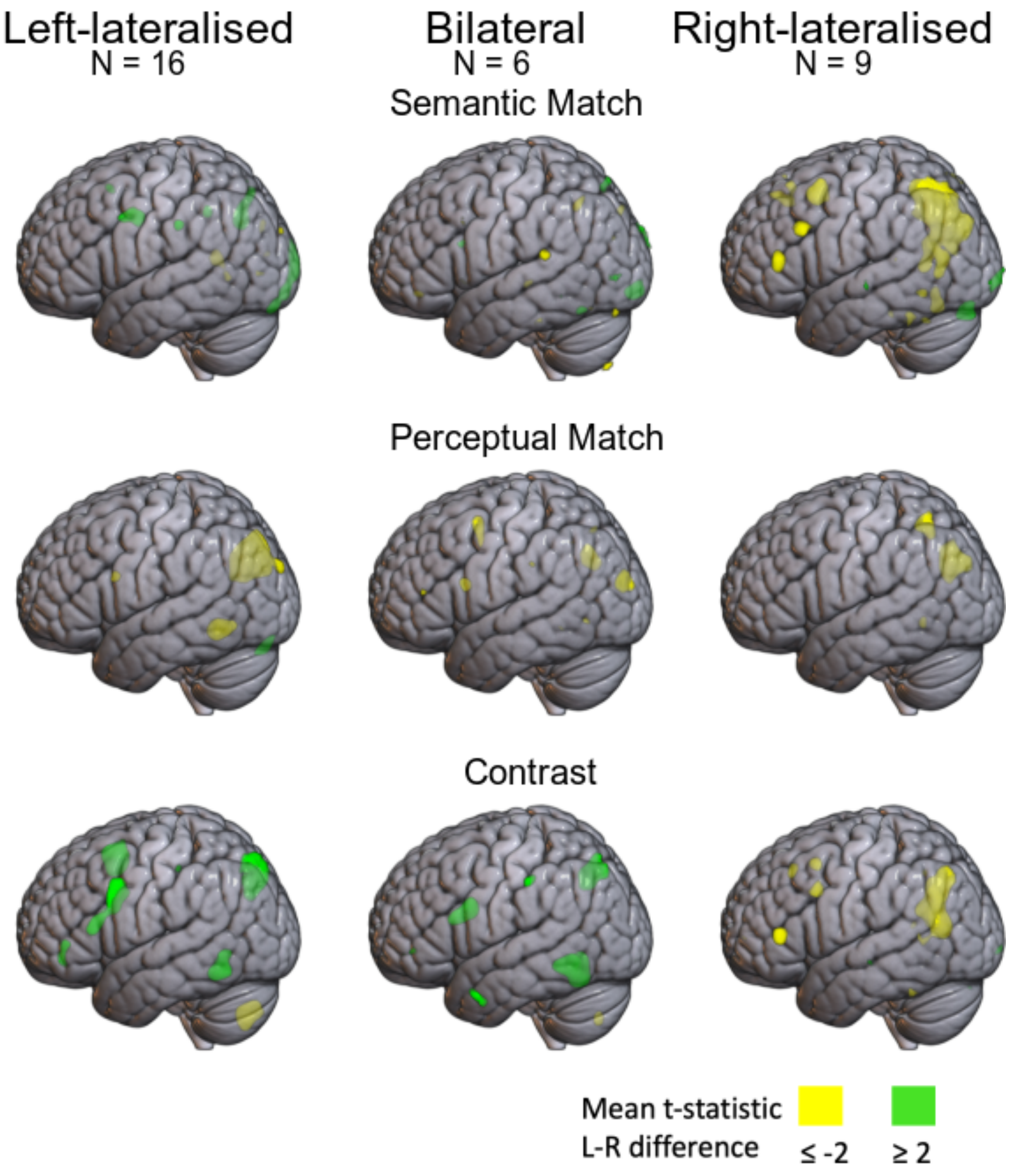


*Figure S3: Left minus right t-statistic maps for the three contrasts in semantic matching task; averages for groups divided according to lateralisation category on fTCD Word Generation.*

Spearman correlations between the Semantic Matching task with no baseline and the Semantic matching contrast with Perceptual matching were 0.73 for the mirror method, and 0.67 for the toolbox method (N = 31).

### Supplementary Material 2

#### Left vs right activations on fMRI by task and region

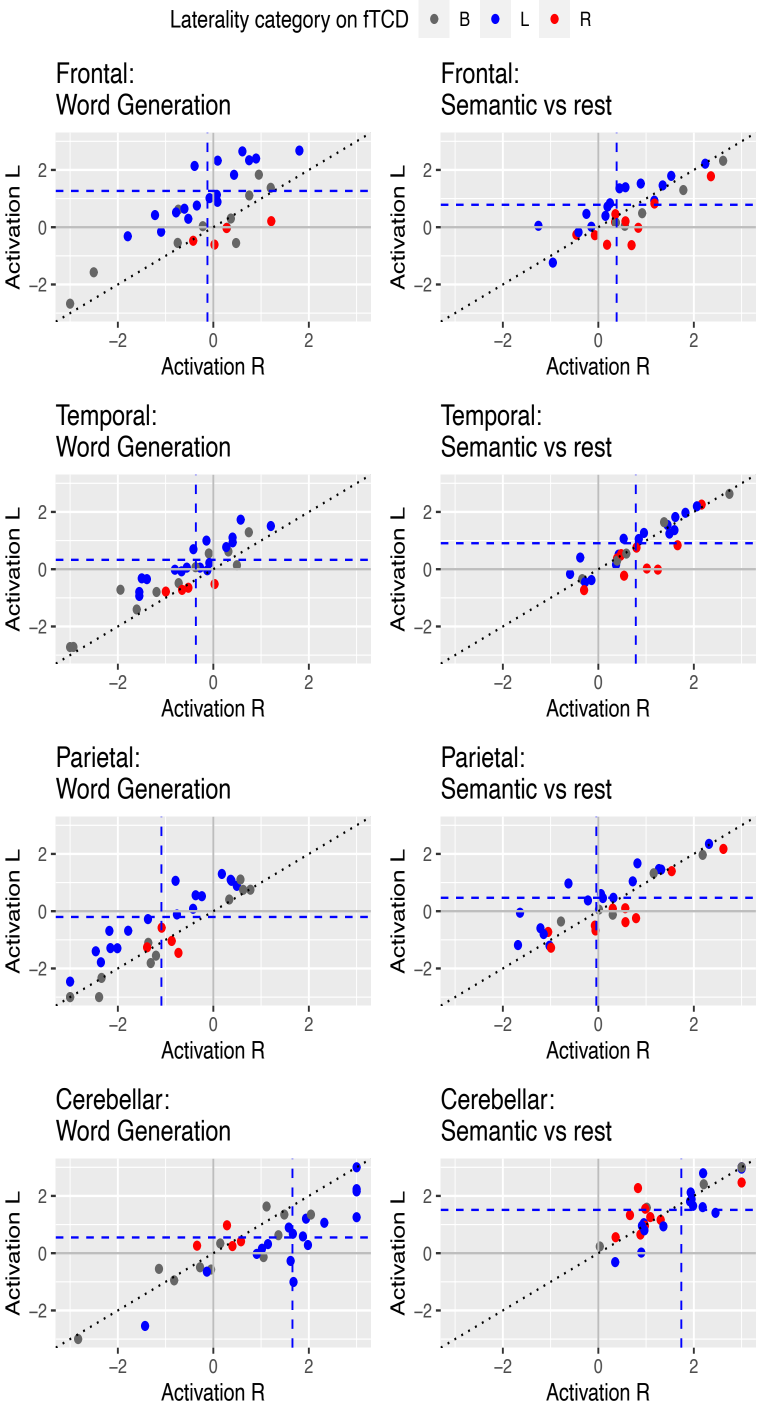


*Figure S4: Mean L and R activations by task and ROI: dotted blue lines show mean for typically (L) lateralised individuals, as categorised on fTCD*

Figure S4 shows the scatterplots with mean activations on left and right. The sloping dotted line is the point of equivalence for left and right: as expected, for the three ROIs contributing to the MCA, individuals who are categorised as left-lateralised tend to fall above this line, indicating more activation on left than right, and those who are right-lateralised tend to fall below the line. For the cerebellar mask, this pattern is reversed, with left-lateralised individuals showing greater activation on the right, and vice versa.

The plots also show different patterns across ROIs in the levels of activation in the two hemispheres. We will focus here just on typically-lateralised individuals, whose means are indicated by the dotted blue lines. The sample size is too small for statistical analysis, but we can form some general impressions on visual inspection. Considering first word generation: in the frontal lobe, typically-lateralised individuals tend to have positive activation in the left hemisphere, and zero activation in the right hemisphere. In the temporal lobe, there is a lesser degree of positive activation in the left hemisphere, coupled with an equivalent deactivation (i.e. mean values below zero) in the right hemsiphere. In the parietal lobe, the mean activation is zero in the left hemisphere, but negative in the right hemisphere. In the cerebellum, there is positive activation in both hemispheres, but greater in the right than the left.

The profile for Semantic matching is rather different, with a less striking difference between the two sides, and no sign of the parietal lobe right hemisphere deactivation seen with Word Generation.
